# Temporal Organization of Synaptic Input and Intrinsic Excitability Shape Direction Selectivity in the Developing *Xenopus laevis* Optic Tectum

**DOI:** 10.64898/2026.09.23.753923

**Authors:** Yugarshi Mondal, Kaiyuan Zheng, Ronald L. Calabrese, Kara G. Pratt

## Abstract

A moving object passes through locations in visual space in a particular order. To istinguish opposite directions of motion, visual circuits must convert this temporal order into different direction selective neuronal responses. This ability emerges and is refined as visual circuits develop. In *Xenopus laevis* tadpoles, direction selectivity in the optic tectum sharpens over development. Companion work by Zheng et al (2026) shows that direction selectivity sharpens between stage 45 (approximately 6–7 days postfertilization), when the retinotectal circuit is undergoing rapid refinement, and stage 48 (approximately 10–16 days postfertilization), when the projection has completed one phase of refinement and visual acuity has improved. The same study shows that, over this period, both the excitatory synaptic input received by tectal neurons and the intrinsic properties of those neurons change. How these synaptic and intrinsic changes work together to produce a sharpening of direction selectivity unclear. Here, we combined visually evoked excitatory postsynaptic current trains recorded at each stage with stage-specific single-compartment conductance-based models. These simulations reproduced the developmental increase in direction selectivity and reduction in spiking at stage 48. Changing intrinsic properties produced only modest changes in spike counts, while reassigning amplitudes among the original event times did not eliminate developmental sharpening. Together, these results suggest that the temporal organization of synaptic input carries much of the directional information, while intrinsic maturation lowers overall spike output, making the input’s existing directional bias more pronounced.

## INTRODUCTION

A moving object passes through locations in visual space in a particular order. To distinguish opposite directions of motion, visual circuits must convert this temporal order into different neuronal responses with a preferred and a null direction. In the retina, asymmetric inhibition suppresses responses to null-direction motion (Barlow & Levick 1965). In visual cortex, differently timed synaptic inputs can be combined approximately linearly to produce a direction-biased membrane-potential response (Jagadeesh et al 1993, Jagadeesh et al 1997). The spike-threshold nonlinearity can then amplify this subthreshold directional bias (Jagadeesh et al 1997, Priebe & Ferster 2005). In each case, synaptic input carries a direction-dependent signal, while the postsynaptic neuron determines how that signal is expressed as spikes.

As visual circuits mature, neurons can become increasingly selective for direction of motion, improving an organism’s ability to distinguish between directions. In the developing *Xenopus laevis* optic tectum, stage 45 occurs during a period of rapid retinotectal refinement, whereas by stage 48 the projection has completed one phase of refinement and visual acuity has improved (Liu et al 2016, Pratt & Aizenman 2007, Sakaguchi & Murphey 1985). Companion work by Zheng et al (2026) found that direction selectivity sharpens between these stages alongside changes in the synaptic input and a decrease in intrinsic excitability. Because these changes were characterized at the population level, their combined consequences for direction-selective spiking could not be inferred directly. It therefore remains unclear whether they are sufficient, when combined, to reproduce developmental sharpening and how each factor contributed.

Here we examined this problem by using reconstructed synaptic inputs from the times and amplitudes of visually evoked excitatory events (EPSCs) recorded at stages 45 and 48 and delivered them to conductance-based neuron models representing neurons at each stage. First we reproduced the developmental sharpening observed by Zheng et al (2026). We then changed the intrinsic properties of the model neurons while holding each input fixed, allowing us to isolate the effect of intrinsic maturation. Finally, we reassigned amplitudes among the original event times to disrupt the association between event amplitude and the preceding interval while preserving event count and temporal organization. Changing intrinsic properties produced only modest changes in spike counts, while amplitude reassignment did not eliminate developmental sharpening. Together, these results suggest that developmental sharpening of direction selectivity arises from the combination of directionally structured synaptic input and reduced intrinsic excitability. In particular, the temporal organization carries much of the directional information and the developmental reduction in intrinsic excitability then amplifies this bias.

## RESULTS

### Stage-specific synaptic inputs and intrinsic properties reproduce developmental sharpening

Zheng et al (2026) found that direction selectivity sharpens between stages 45 and 48 alongside decreases in both the overall excitatory synaptic drive received by tectal neurons and their intrinsic excitability, with the latter driven primarily by reduced maximal sodium conductance. We asked whether these coordinated synaptic and intrinsic changes were sufficient to reproduce the sharpening of spike output. As described in Methods, we reconstructed visually evoked excitatory inputs from the times and amplitudes of recorded synaptic events and delivered them to model neurons with the intrinsic properties of the corresponding developmental stage. Example simulations produced more spikes in the preferred direction than in the null direction at both stages (Fig. 1A). Across inputs, the stage 45 input–cell pairing produced median spike counts of 33 in the null direction and 46 in the preferred direction, with a DSI of 0.283. The stage 48 pairing produced fewer spikes overall—median counts of 7 and 12, respectively—but a higher DSI of 0.458 (Fig. 1B). Thus, the stage-specific synaptic inputs and intrinsic properties were sufficient, when combined in a model, to reproduce increased direction selectivity despite reduced overall spiking.

**Figure 1.**
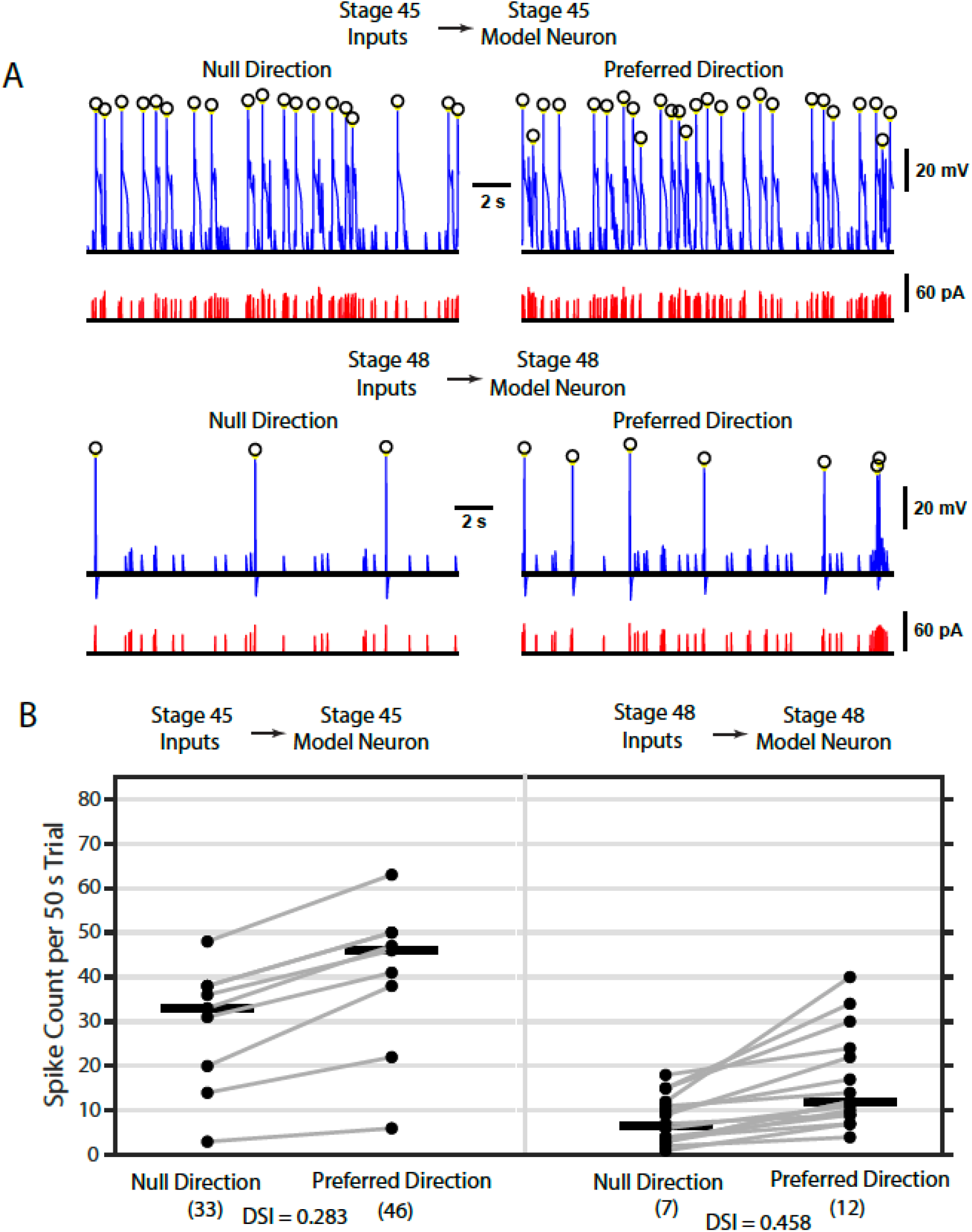
Combining synaptic inputs and intrinsic properties from each stage reproduces developmental sharpening of direction selectivity. (A) Representative null- and preferred-direction simulations using stage 45 inputs and the stage 45 model neuron (top) or stage 48 inputs and the stage 48 model neuron (bottom). Blue traces show membrane voltage, red traces show the reconstructed inward synaptic current, and circles mark detected spikes. The traces show excerpts from the simulated trials. (B) Spike counts over each 50-s trial for stage 45 inputs delivered to the stage 45 model neuron (left) and stage 48 inputs delivered to the stage 48 model neuron (right). Each dot represents one direction-specific trial, gray lines connect paired null- and preferred-direction trials, and black bars show median counts. Numbers beneath the direction labels are rounded medians. Direction-selectivity index (DSI) was calculated from the unrounded medians as 1-ND/PD, where ND and PD are the median null- and preferred-direction spike counts. The analysis included 9 stage 45 and 16 stage 48 trial pairs. Pairs in which either direction produced no spikes were excluded, and pairs with DSI greater than 0.20 were retained.

### Intrinsic maturation produces modest changes in preferred and null-direction spike counts

We sought to isolate the contribution of intrinsic maturation. We first delivered stage 45 inputs to model neurons with either stage 45 or stage 48 intrinsic properties. When these inputs were delivered to the stage 45 versus the stage 48 model neuron, median null-direction spike counts decreased from 33 to 31, while median preferred-direction counts increased from 46 to 47. (Fig. 2A). We next delivered stage 48 inputs to both model neurons. When these inputs were delivered to the stage 45 model versus the stage 48 model neuron, median null-direction counts decreased from 10 to 7 and median preferred-direction counts decreased from 15 to 12 (Fig. 2B). Thus, stage 45 inputs continued to produce high spike counts in the stage 48 model neuron, whereas stage 48 inputs continued to produce low spike counts in the stage 45 model neuron. This suggests the synaptic input largely determined whether the model produced a high- or low-spiking response. Moreover, while intrinsic maturation had only modest effects on absolute spike counts it still could increase DSI, especially when the synaptic input placed the neuron in a low-spiking regime.

**Figure 2.**
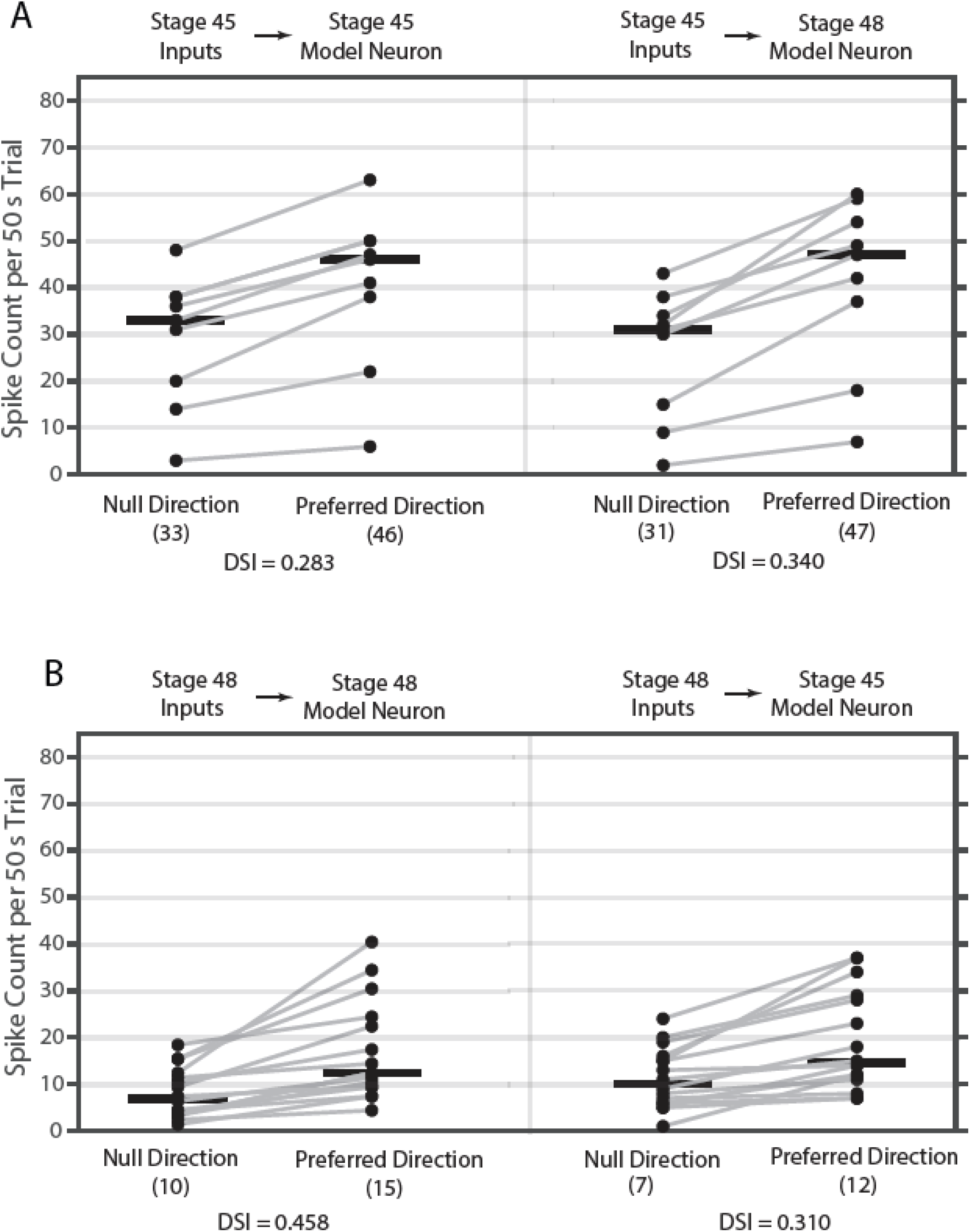
Intrinsic maturation leaves absolute spike counts largely unchanged while increasing direction selectivity. (A) The nine stage 45 input pairs shown in Figure 1B were delivered to the stage 45 model neuron (left) or the stage 48 model neuron (right). The left condition repeats the stage 45 results from Figure 1B. (B) The 16 stage 48 input pairs shown in Figure 1B were delivered to the stage 48 model neuron (left) or the stage 45 model neuron (right). The left condition repeats the stage 48 results from Figure 1B. Plotting conventions and DSI calculations are as in Figure 1B.

### Developmental sharpening persists after event amplitudes are reassigned

In the pooled biological data from Zheng et al (2026), events following shorter interevent intervals tended to have larger amplitudes. To incorporate this association, we assigned each modeled event an amplitude based on its preceding interevent interval, with shorter intervals producing larger events (see Methods). We therefore first compared the IEI distributions across developmental stages and directions. Preferred-direction inputs contained shorter intervals than null-direction inputs at both stages, while stage 45 inputs contained shorter intervals overall than stage 48 inputs (Fig. 3B). As expected from the amplitude-assignment rule, these timing differences were mirrored in the amplitude distributions: preferred-direction events were larger overall than null-direction events, and stage 45 events were larger overall than stage 48 events (Fig. 3A).

**Figure 3.**
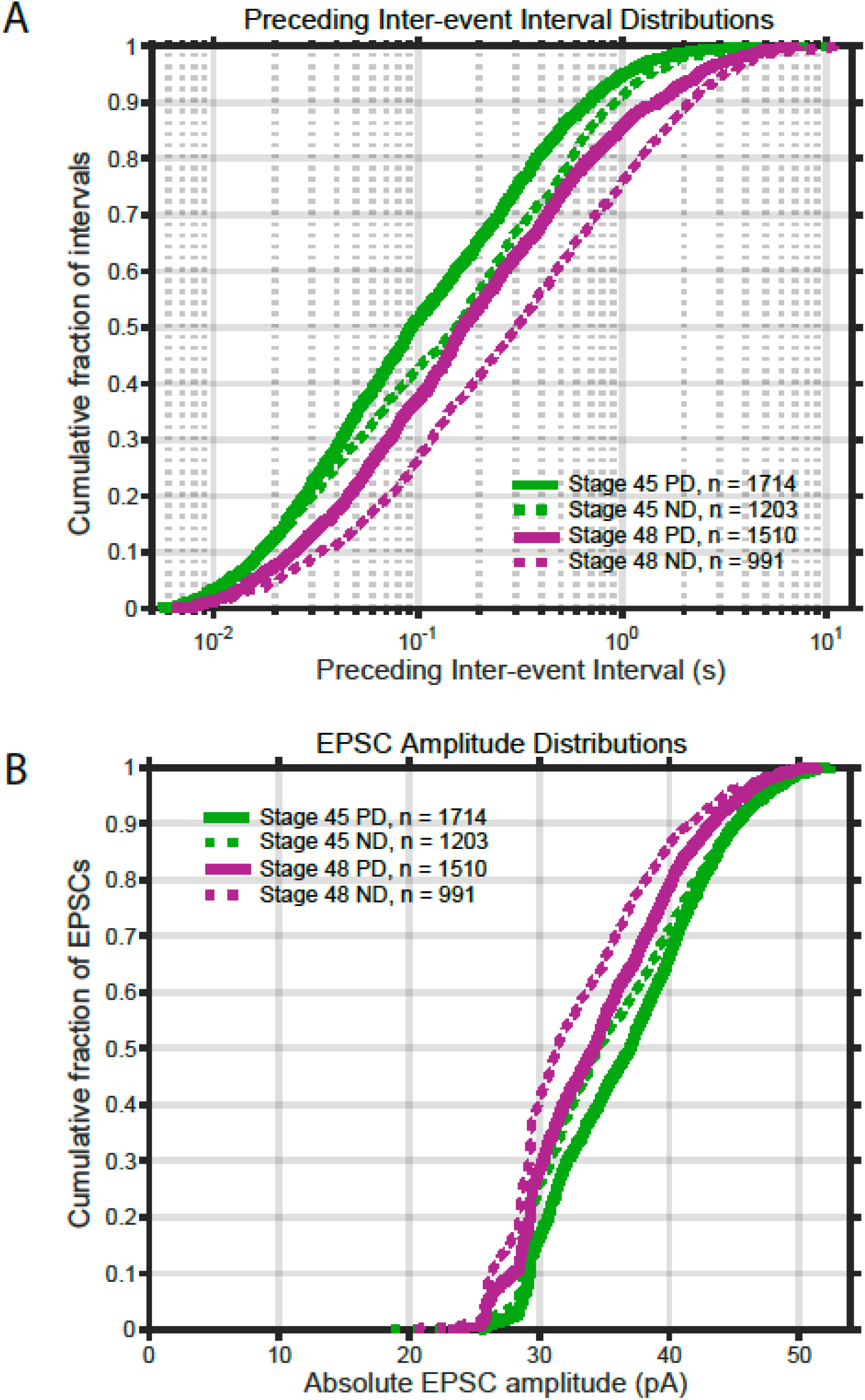
Distributions of interevent intervals and amplitude in the reconstructed synaptic inputs. (A) Empirical cumulative distributions of the preceding inter-event intervals for stage 45 and stage 48 preferred-direction and null-direction inputs. The horizontal axis shows the interval preceding each event. (B) Empirical cumulative distributions of the absolute amplitudes assigned to those events. Stage 45 inputs are shown in green and stage 48 inputs in purple. Solid lines indicate preferred-direction inputs and dashed lines indicate null-direction inputs. *n* indicates the number of events.

So, we asked whether direction-selective output depended on this specific assignment of amplitudes to event times. Within each trial, we kept the number and timing of the events fixed but randomly reassigned the recorded amplitudes among those times. This manipulation preserved the event count, temporal pattern, and complete set of amplitudes within each trial while disrupting the original association between amplitude and timing.

Preferred-direction inputs continued to produce more spikes than null-direction inputs after amplitude reassignment (Fig. 4A). In a second reassignment, the stage 45 input–cell pairing produced median null- and preferred-direction counts of 50 and 63 spikes, respectively, with a DSI of 0.206. The stage 48 pairing produced median counts of 8 and 17 spikes, with a DSI of 0.515 (Fig. 4B). Across all five independent amplitude reassignments, DSI remained higher for the stage 48 pairing than for the stage 45 pairing (Fig. 4C).

**Figure 4.**
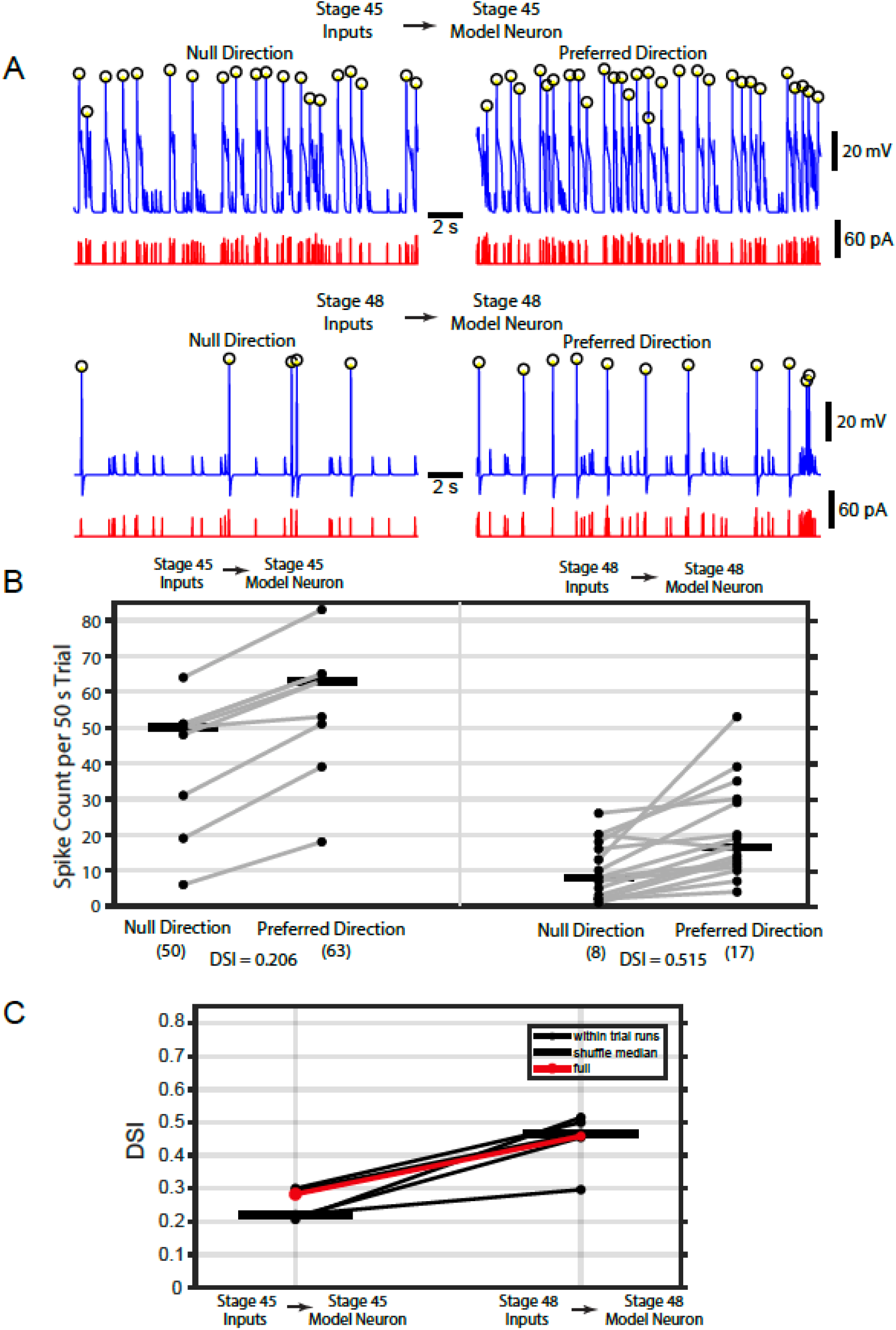
Developmental sharpening persists after amplitudes are reassigned among event times. Within each 50-s trial, event amplitudes were randomly reassigned among the original event times. (A) Representative voltage traces (blue) and reconstructed synaptic currents (red) from the first reassignment. Following Figure 1A, circles mark detected spikes; the traces show excerpts from the simulated trials. (B) Paired null- and preferred-direction spike counts from all pairs in Figure 2. (C) DSI after amplitudes were randomly redistributed five separate times. Each black line shows the results of one complete rerun, the thick black bars show the median across the five reruns, and the red line shows the original simulations from Figure 2 without amplitude redistribution.

Although amplitude reassignment altered the precise spike counts and DSI, it did not eliminate developmental sharpening. Because event times were unchanged, clusters of closely spaced events remained intact while the amplitudes assigned to events within those clusters were rearranged. This seems to suggest an important role for temporal clustering. Directly testing this, however, would require manipulating event timing.

## DISCUSSION

The present results separate the contributions of synaptic and intrinsic development. Synaptic inputs carried a directional bias at both developmental stages: preferred-direction inputs elicited more spikes than their paired null-direction inputs in either model neuron (Fig. 1B, 2A, 2B). Switching between the stage 45 and stage 48 model neurons altered absolute spike counts only modestly (Fig. 2A, 2B). Thus, in the model, the developmental difference in spike counts was driven primarily by the stage-specific synaptic inputs rather than by intrinsic maturation. Nevertheless, the modest changes produced by intrinsic maturation substantially affected DSI, especially for stage 48 inputs. For the same stage 48 inputs, replacing the stage 45 model neuron with the less excitable stage 48 model neuron reduced the median preferred and null responses by three spikes, from 15 to 12 and from 10 to 7, respectively (Fig. 2B). These equal absolute reductions had unequal proportional effects: 20% of the preferred response but 30% of the null response. Because DSI = 1 − ND/PD, the resulting decrease in ND/PD from 10/15 to 7/12 increased DSI even though the absolute five-spike difference remained unchanged. In the low-spiking regime produced by stage 48 inputs, each spike had greater influence on the preferred-to-null response ratio, allowing a small absolute change in spike count to produce a substantial change in DSI. Intrinsic maturation therefore did not create the directional preference; it made the existing synaptic bias more consequential in the low-spiking stage 48 regime.

Furthermore, developmental sharpening of direction selectivity persisted after amplitudes were reassigned among the original event times. This manipulation changed which amplitudes occurred within clusters of closely spaced events but left the clusters themselves intact. The persistence of sharpening therefore suggests that temporal clustering contributes to the directional bias independently of the precise placement of individual event amplitudes.

Together, these simulations show that synaptic input supplies the directional bias, while cellular properties determine how strongly that bias is expressed in spike output. Our simulations reproduced the developmental increase in DSI observed by Zheng et al (2026), and both studies showed a developmental decrease in null-direction firing. The agreement did not extend fully to preferred-direction firing: Zheng et al (2026) did not observe a developmental difference, whereas we found that preferred-direction spike counts were lower in the stage 48 input–cell simulations than in the stage 45 simulations. This discrepancy may partly reflect how the experimental and simulated populations were constructed. Zheng et al (2026) compared separate populations of stage 45 and stage 48 neurons, preserving the natural cell-to-cell variation within each stage. In our simulations, input trains from both stages were reconstructed and adjusted using a common procedure and then delivered to a common model neuron for each stage. The resting membrane potential and other intrinsic properties of the neuron from which each input was recorded were therefore replaced by those of the corresponding model neuron. This standardization allowed us to separate the effects of synaptic input stage and cellular (i.e. intrinsic excitability; output) stage, but it removed much of the neuronal variability present in the experimental populations. Differences between the experimental and simulated preferred- and null-direction responses may therefore reflect this standardization as well as other simplifications of the model.

Prior work suggests that synaptic input and neuronal output interact during development of the *Xenopus* optic tectum. Changes in synaptic drive regulate intrinsic excitability, which shapes neuronal spike output (Pratt & Aizenman 2007). In turn, chronically reducing tectal spike output disrupts receptive-field refinement, showing that neuronal output can influence subsequent circuit development (Dong & Aizenman 2012). Together, these findings suggest that synaptic input can alter neuronal output and that neuronal output can influence subsequent circuit refinement. Zheng et al (2026)found that synaptic input and intrinsic excitability both change between stages 45 and 48 as direction selectivity sharpens. Our simulations separate the contributions of these changes to spike output. However, they cannot determine whether feedback between synaptic input, intrinsic excitability, and circuit refinement produced the developmental changes.

Temporal organization may contribute both to the development and to the expression of direction selectivity. Engert et al (2002) showed that that appropriately patterned visual stimuli induced direction sensitivity in developing tectal neurons. This finding indicates that temporally patterned activity can produce direction-selective refinement. In our simulations, developmental sharpening persisted after amplitudes were reassigned while clusters of closely spaced events remained intact, suggesting that temporal clustering also contributes to the expression of directional bias in synaptic input. Together, these findings suggest that the temporal organization of synaptic inputs may come to carry a directional bias that is further refined over development.

Developmental sharpening of direction selectivity occurs across visual systems, but it can arise through different synaptic and intrinsic changes in different species. In developing ferret visual cortex, direction-selectivity sharpening was accompanied by increased intrinsic excitability and an increased subthreshold voltage response to preferred-direction motion (Roy et al 2020). In the developing *Xenopus* optic tectum, sharpening was accompanied by decreased intrinsic excitability and a decreased spiking response to null-direction motion. The shared principle is therefore not a particular direction of change in excitability. Rather, development can sharpen direction selectivity by changing both the synaptic input that establishes a directional bias and the intrinsic properties that govern its nonlinear conversion into spikes.

## METHODS

### The Model

We used a single-compartment Hodgkin-Huxley-type model to represent tectal neurons at stages 45 and 48. The model was designed to examine how stage-specific excitatory input is converted into spike output. It contained a fast sodium current (I_Na_), a delayed-rectifier potassium current (I_Kd_), an A-type potassium current (I_A_), and a leak current (I_L_). These currents were chosen to represent the major inward, sustained outward, and transient outward components observed in voltage-clamp recordings (Zheng et al 2026). Synaptic input was delivered as a current waveform rather than through a conductance-based synapse.

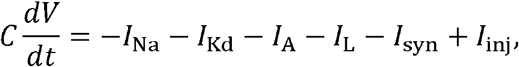

where *I*_syn_ denotes the positive magnitude of inward synaptic current. The intrinsic currents were

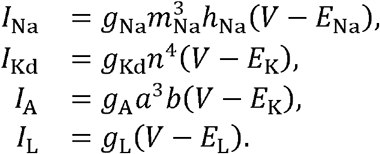

The membrane capacitance (C) was 15 pF. Reversal potentials were *E*_NA_ = 60mV, *E*_k_ = −80mV, and *E*_k_ = −50mV. The shared maximal conductances were *g*_Kd_ = 0.7nS, *g*_A_ = 0.7nS, and *g*_L_ = 0.3nS

The stage-specific models differed only in maximal sodium conductance. The stage 45 model used *g*_*Na*_ = 11nS, whereas the stage 48 model used *g*_*Na*_ = 5nS. This reduction represented the developmental decrease in sodium current identified in the accompanying voltage-clamp measurements (Zheng et al 2026). All other conductances and channel kinetics were held constant. Thus, within the present model, intrinsic maturation was represented specifically by a reduction in maximal sodium conductance.

Each gating variable x followed

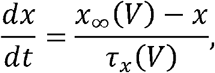

Where

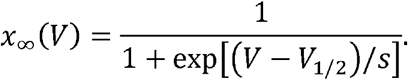

The half-activation or half-inactivation voltages and slope factors were (−32, −5) mV for sodium activation, (−50, −5) mV for sodium inactivation, (−25, −5) mV for delayed−rectifier activation, (−38, −5) mV for A-current activation, and (−50, −5) mV for A-current inactivation. Voltage-dependent time constants, in milliseconds, were

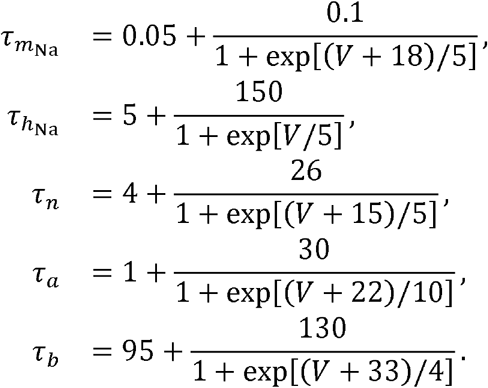

Simulations began at −50 mV, with all gating variables initialized to their steady-state values at that voltage.

### Voltage-clamp and current-ramp characterization of the model neurons

The intrinsic-current models were evaluated using the voltage-command waveforms applied in the experimental voltage-clamp recordings. Ten command traces were read from the experimental voltage-step file obtained from Zheng et al (2026) and imposed on both model neurons. Under voltage clamp, membrane voltage was fixed by the command waveform while the channel-gating equations were integrated. The sodium, delayed-rectifier, A-type, and leak currents were then calculated and summed to obtain the predicted whole-cell ionic current. The maximal conductances were selected by comparing these simulated current families with the stage-specific experimental recordings rather than through an automated fit (Fig. 5A).

**Figure 5.**
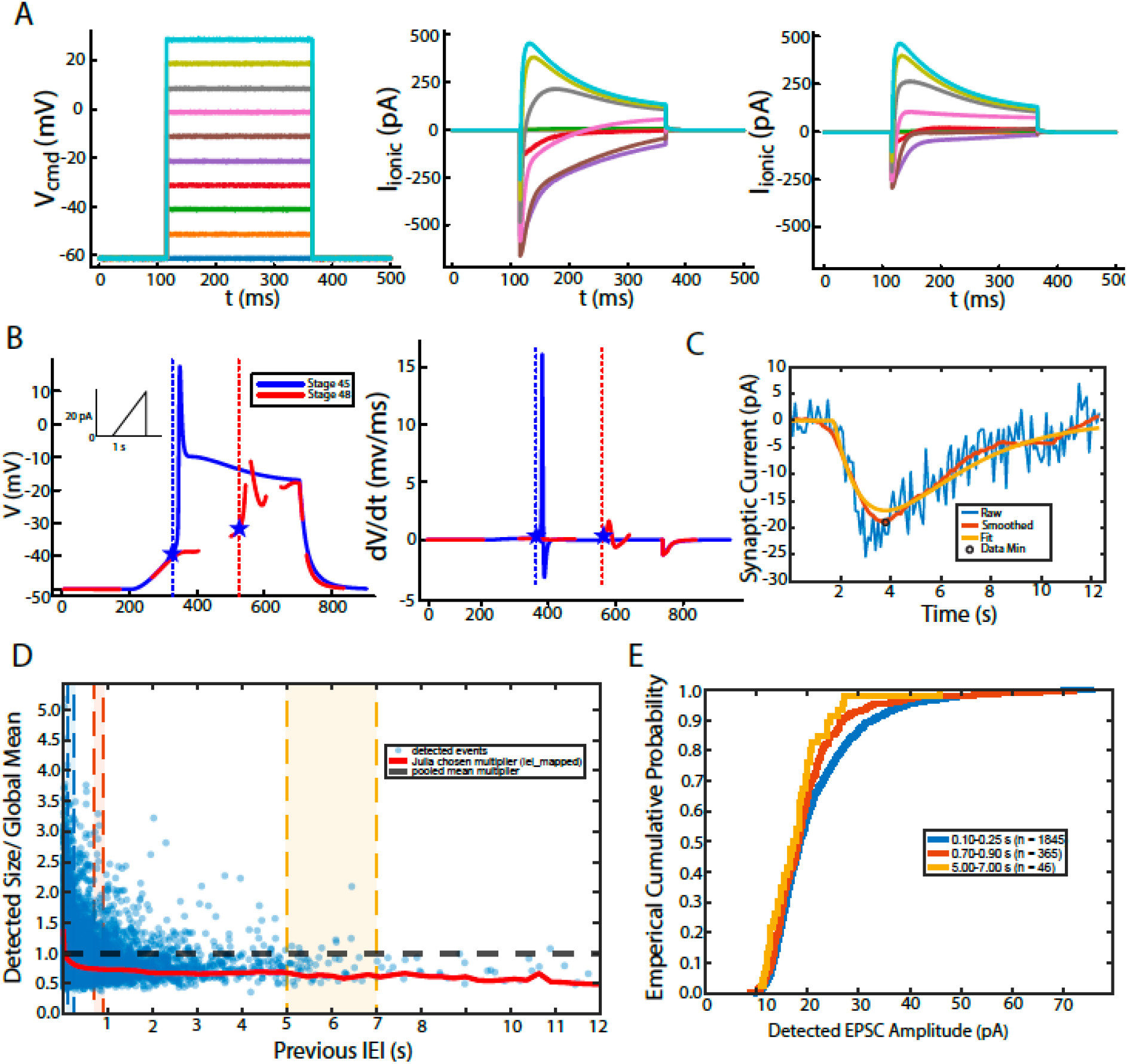
Construction of the stage-specific neuron models and simulated synaptic inputs. (A) Voltage-command family used to characterize the intrinsic-current models (left) and the resulting current response in the stage 45 (middle) and stage 48 (right) model neurons. Colors identify corresponding voltage commands and current responses. The models differed only in maximal sodium conductance, which was 11 nS in the stage 45 model and 5 nS in the stage 48 model. (B) Voltage responses (left) and their time derivatives (right) during the same current ramp in the stage 45 (blue) and stage 48 (red) model neurons. The inset shows the ramp from 0 to 20 pA over 500 ms. Stars mark action-potential onset, and dotted lines indicate the corresponding onset times. (C) Representative fit of the simulated EPSC waveform to an isolated synaptic event. The raw current trace is shown in blue, the trace after applying a 2-ms moving average is shown in red, and the fitted double exponential is shown in yellow. The open circle marks the minimum of the smoothed trace used to align the fit. (D) Relationship between detected EPSC amplitude and the preceding interevent interval. Each blue point represents one detected event, with its amplitude divided by the mean amplitude of all pooled events. The black dashed line marks the pooled mean. The red line shows the relationship between the amplitude multiplier and the preceding interval used in the simulation. Colored vertical bands mark the three representative interval ranges examined in E. (E) Cumulative distributions of detected EPSC amplitudes for interevent intervals of 0.10–0.25 s, 0.70–0.90 s, and 5.00–7.00 s. Sample sizes indicate the number of events in each interval range. The amplitude-assignment procedure imposed the weak association observed in the recordings between shorter preceding intervals and larger events.

We also selected the conductances so that the two models had distinct spike thresholds, simulating the difference seen in Zheng et al (2026). A common current ramp began at 200 ms and increased linearly from 0 to 20 pA over 500 ms. We calculated *dv/dt* using centered differences. Action-potential onset was defined as the first point before the initial spike peak at which *dv/dt* exceeded the mean plus five standard deviations of *dv/dt* during the first 50 ms of the ramp. This procedure produced thresholds of −39.32 mV for the stage 45 model and −32.04 mV for the stage 48 model (Fig. 5B).

### Experimental synaptic-event data and selection of input streams

Visually evoked excitatory synaptic events were obtained from the stage 45 and stage 48 recordings described by Zheng et al. (2026). In that study, synaptic-input and spike-output direction selectivity were measured in separate populations of tectal neurons. Motion-evoked EPSCs were recorded under voltage clamp, whereas spike responses were recorded from different neurons in loose cell-attached mode; input and output selectivity were therefore not paired within individual cells. Synaptic-event summary files from the voltage-clamp recordings were used in the present study. For each event, these files contained the preceding interevent interval, detected current amplitude, trial identity, stimulus direction, and direction-selectivity measurement associated with the recording. No new electrophysiological recordings were performed for the present study.

The left–right and up–down direction pairs were analyzed separately. A pair was included if the DSI calculated from the magnitudes of the recorded responses exceeded 0.20. Within each qualifying pair, the direction producing the larger response was designated the preferred direction and the opposite direction the null direction.

Preferred- and null-direction trials were paired according to the trial assignments in the experimental summary files. Each trial was reconstructed independently over a fixed duration of 50 s. Starting at time zero, the supplied interevent intervals (IEIs) were cumulatively summed to determine successive event times. An event was retained if its reconstructed time was less than or equal to 50 s.

### Estimation of the synaptic-current kernel

The waveform of an individual simulated EPSC was measured from recordings containing isolated synaptic events, also obtained from Zheng et al (2026). For each recording, the mean current during the first 1 ms was subtracted from the entire trace, setting the pre-event baseline to approximately zero. Current was then converted from amperes to nanoamperes. A 2-ms moving average was applied for fitting (Fig. 5C).

Because the onset time of each isolated EPSC could not be identified reliably, each waveform was aligned to its most negative point. The minimum of the smoothed current trace was identified and treated as the EPSC peak. The waveform from 5 ms before to 200 ms after this minimum was fit by nonlinear least squares with a negative-going double exponential:

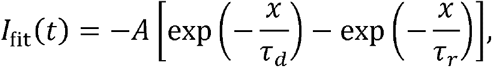

Where

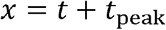

and

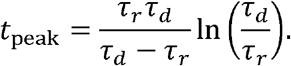

Here, *t*_*peak*_ is the onset-to-peak interval implied by the candidate rise and decay constants. The transformation *x* = *t* + *t*_*peak*_ converts the peak-centered time coordinate of the recorded waveform into time since the inferred EPSC onset. Consequently, the model onset occurs at *t =* − *t*_*peak*_, and its analytical peak occurs at the observed current minimum, *t* = 0. The fitted current was set to zero before the inferred onset.

Initial estimates of *τ*_*r*_ and *τ*_*d*_ were obtained from the times at which the waveform crossed 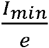 before and after the observed peak, respectively. These estimates were used only to initialize the nonlinear fit. The fitted rise constant was constrained to 0.05–10 ms, and the decay constant was constrained to 0.2–15,000 ms.

A total of 52 isolated EPSCs produced finite, positive parameter estimates. The median fitted rise and decay constants were 1.670 and 2.369 ms, respectively. Every simulated EPSC used these median time constants, whereas EPSC amplitudes were assigned using the empirical procedure described below.

### The relationship between event amplitude and interevent interval

Event timing was taken from the recorded IEIs, but the amplitude measured for each particular event was not transferred directly to the corresponding event in the simulated train. Instead, amplitudes were assigned using the empirical relationship between detected EPSC amplitude and the preceding IEI.

To construct this relationship, detected EPSCs from the selected stage 45 and stage 48 preferred- and null-direction trials were pooled. Current amplitudes were converted to positive magnitudes and divided by the mean amplitude of all retained events to obtain dimensionless amplitude multipliers, *M*_*i*_. Events with preceding IEIs between 0.001 and 12 s were used to construct an IEI-conditioned empirical inverse cumulative distribution.

The IEI range was divided into 45 linearly spaced centers. Around each center, a symmetric window was initially assigned a half-width of 0.12 s. The window was expanded in increments of 25% until it contained at least 150 events or reached a maximum half-width of 0.75 s. Within each IEI window, the amplitude multipliers were sorted from smallest to largest, and the multiplier corresponding to each quantile level from 0.001 to 0.999 was computed. These values described the distribution of amplitude multipliers observed for events with preceding IEIs in that range. Repeating this procedure at all 45 centers described how the amplitude distribution changed with preceding IEI (Fig. 5D).

The preceding calculations produced empirical amplitude distributions for different ranges of preceding IEIs (Fig. 5E). We used these distributions to map each simulated event’s preceding IEI to an amplitude multiplier. Loosely speaking, shorter IEIs were assigned amplitudes from higher percentiles of the corresponding distribution, whereas longer IEIs were assigned amplitudes from lower percentiles. The mapping was defined as follows.

IEIs were first restricted to the range 0.005–12 s and transformed onto a normalized logarithmic scale:

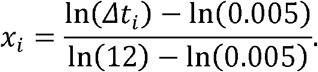

For each event, the preceding IEI is assigned a quantile. Shorter IEIs selected higher quantiles, as defined by:

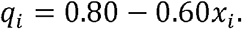

Thus, the shortest intervals were assigned the 80th-percentile amplitude, whereas the longest intervals were assigned the 20th-percentile amplitude. The selected quantile was used to obtain the corresponding amplitude multiplier from the empirical distribution. This multiplier was then converted into an event amplitude according to:

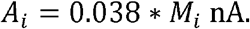

The scale factor of 0.38 nA was a model parameter chosen so that null-direction inputs could evoke spikes while preferred- and null-direction responses remained distinguishable. The relationship between preceding IEI and assigned amplitude is shown by the red line in Fig. 5D.

By construction, shorter preceding intervals tended to receive larger event amplitudes. This association should not be interpreted as evidence for a particular physiological mechanism, such as synaptic facilitation. Rather, the input-construction procedure enforced the weak trend observed in the recordings, in which shorter preceding IEIs tended to be associated with larger EPSCs (Zheng et al 2026).

### Construction of synaptic-current trains

Each assigned event was converted into a peak-normalized double-exponential current:

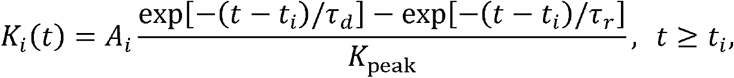

where *t*_*i*_ was the reconstructed event time and *K*_*peak*_ was the maximum of the unnormalized double exponential. This normalization made *A*_*i*_, from above, the peak inward-current magnitude of the simulated event. The total synaptic current was obtained by summing:

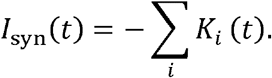

Every event used *τ*_*r*_ = 1.670ms and *τ*_*d*_ = 2.369 Each kernel was evaluated until 10*τ*_*d*_ after its onset, after which its remaining tail was truncated for computational efficiency. Synaptic-current traces were constructed at 0.1-ms resolution.

### Numerical implementation

Model simulations were implemented in Julia using DifferentialEquations.jl. Differential equations were integrated with the Tsitouras 5/4 Runge-Kutta method. Voltage and state variables were saved every 0.1 ms, with a maximum internal step of 0.5 ms, a relative tolerance of 10^−6^, and an absolute tolerance of 10^−9^ Experimental-data preprocessing, EPSC fitting, output analysis, and figure generation were performed in MATLAB.

## DATA AVAILABILITY

The code and parameters required to reproduce the results in this work are accessible at Zenodo.

## ACKNOWLEDGMENTS

YM acknowledges the support of NIMH (R01MH046742) and the fruitful scientific conversations with Eve Marder. This work was also supported by the National Science Foundation (2212591) to KGP.

## REFERENCES

Barlow HB, Levick WR. 1965. The mechanism of directionally selective units in rabbit’s retina. J Physiol 178: 477–504

Dong W, Aizenman CD. 2012. A competition-based mechanism mediates developmental refinement of tectal neuron receptive fields. J Neurosci 32: 16872–9

Engert F, Tao HW, Zhang LI, Poo MM. 2002. Moving visual stimuli rapidly induce direction sensitivity of developing tectal neurons. Nature 419: 470–5

Jagadeesh B, Wheat HS, Ferster D. 1993. Linearity of summation of synaptic potentials underlying direction selectivity in simple cells of the cat visual cortex. Science 262: 1901–4

Jagadeesh B, Wheat HS, Kontsevich LL, Tyler CW, Ferster D. 1997. Direction selectivity of synaptic potentials in simple cells of the cat visual cortex. J Neurophysiol 78: 2772–89

Liu Z, Hamodi AS, Pratt KG. 2016. Early development and function of the Xenopus tadpole retinotectal circuit. Curr Opin Neurobiol 41: 17–23

Pratt KG, Aizenman CD. 2007. Homeostatic regulation of intrinsic excitability and synaptic transmission in a developing visual circuit. J Neurosci 27: 8268–77

Priebe NJ, Ferster D. 2005. Direction selectivity of excitation and inhibition in simple cells of the cat primary visual cortex. Neuron 45: 133–45

Roy A, Osik JJ, Meschede-Krasa B, Alford WT, Leman DP, Van Hooser SD. 2020. Synaptic and intrinsic mechanisms underlying development of cortical direction selectivity. Elife 9

Sakaguchi DS, Murphey RK. 1985. Map formation in the developing Xenopus retinotectal system: an examination of ganglion cell terminal arborizations. J Neurosci 5: 3228–45

Zheng K, Mondal Y, Roth B, Udoh UG, Calabrese RL, Pratt KG. 2026. Development of direction selectivity in the Xenopus tadpole optic tectum and midbrain tegmentum. preprint

